# Multibody kinematic optimisation vs body fat: A performance analysis

**DOI:** 10.1101/2022.07.26.501536

**Authors:** Vignesh Radhakrishnan, Samadhan B Patil, Adar Pelah

## Abstract

We have analysed the performance of mulitbody kinematic optimisation methods in reducing soft tissue artefacts for subject data of varying body fat percentages. Multibody kinematic optimisation methods are a critical aspect of movement analysis using musculoskeletal modelling software. By minimising soft tissue artefacts, they help in achieving higher fidelity joint kinematics and dynamics analyses. Prior studies have not examined the performance of multibody kinematic optimisation on subjects of varying body fat percentages. Herein, we: 1) have analysed the efficacy of three different multibody kinematic optimisation methods on varying body fat percentages, 2) implemented a novel weighting scheme to reduce error irrespective of body fat percentages. Residual error using gait data of 50 participants of varying body fat percentages was calculated through inverse kinematic analysis using OpenSim(c) musculoskeletal modelling software. The analysis was repeated using a time-based weighting scheme. The residual error of participants with higher body fat percentages was greater by 30% when compared to residual error of participants of lower body fat percentages. Additionally, time-based weighting scheme reduced residual error by 20% on average compared to constant-value weighting scheme. Our results indicate that multibody kinematic optimisation methods are adversely affected by higher body fat percentages and that time-based weighting can provide higher fidelity movement analysis irrespective of body fat percentages. Through our results we aim to develop tools which provide greater precision in obesity-related movement analysis. Such tools could also help address the disparities in rates of obesity associated with different ethnic or socioeconomic background.

## Introduction

Movement analysis is a widely used clinical tool in the diagnosis, rehabilitation and surgical planning for neurological and musculoskeletal pathologies, with additional applications in sports medicine. [1, 2] Through the use of skin-mounted marker motion capture systems and force plates, kinematic and kinetic values during different activities are calculated to study joints forces, angles and moments. [2, 3]

Skin-mounted marker motion capture systems are clinical gold standards in movement analysis, with sub-millimeter accuracy enabling precise study of movements. The main sources of error in these systems are soft tissue artefacts (STA) and errors caused by marker misplacement. STA are discrepancies in the movement of markers with respect to the underlying bone, caused by inertial effects, skin sliding and deformation, gravity, and muscle contraction. [4, 5] While the latter source of error can be minimised by experienced clinicians, STA are very difficult to compensate for by simple filtering techniques as they possess similar frequencies to that of bone movement, and are task and subject dependent. The effect of soft tissue artefacts on the estimation of joint angles and rotation is substantial, with errors of up to 30mm and 20 degree [6, 7] leading to erroneous estimation of muscle forces [8] and functional joint centres [9] which affects clinical interpretability of data and in some instances invalidating clinical analysis. These factors make them the most critical source of error in skin-mounted marker based systems. [4, 10]

Numerous methods to analyse and compensate for soft tissue artefacts have been proposed over the years. [6] These methods can be categorised into imaging or hardware based and analytical methods. Imaging or hardware based methods such as intracortical or bone-mounted pins, percutaneous markers, X-ray, fluroscopy and MRI have been used to quantify STA during different forms of motions. [4, 6, 11, 12] Analytical methods to compensate for STA can be split into segmental-based methods and multibody kinematic optimisation (MKO) methods. Segmental-based methods such as the solidification technique [13], pliant surface modelling, point cluster technique, and double calibration [4, 14] have been proposed to reduce the effect of deformation of the skin caused during various movements. [6] These segment-based compensation techniques work by treating each segment independently, without imposing joint constraint, which result in non-physiological joint translations and rotations caused by differing skin movement patterns in adjacent segments. [6, 15]

MKO methods are global optimisation methods that have been proposed to overcome the shortcoming of segmental-based approaches i.e of producing non-physiological movements, and are a key aspect of musculoskeletal modelling. By incorporating all segments of the body in a multi-link model with constraints, MKO methods compensate for both, the deformation of the surface marker cluster and the rigid displacement of the cluster. [16] MKO methods, such as the global optimisation method (GOM) [15], Local Marker Estimation (LME) [17, 18] and Kalman Smoothing (KS) [19] reduce soft tissue artefacts by determining the optimal pose of a virtual model in musculoskeletal modelling software. The optimal pose is determined by minimising the error between virtual markers in the modelling software and experimental markers worn by the patient.

Whilst MKO methods have been shown to reduce the influence of soft tissue artefacts on calculated joint kinematics, validation has only been performed on either simulated data, or on data from subjects with a body mass index (BMI) of less than 25, which is considered as a healthy weight for their height. [7, 20] Furthermore, studies which have focused on the effect of subcutaneous adipose on soft tissue artefacts have focused on analysing the effectiveness of new marker sets and marker weights on reducing the influence of soft tissue artefacts, and have not analysed the overall efficacy of the MKO method used. [21–23] Obesity-specific marker sets, which use a combination of traditional markers and digitally probed and placed markers, have been shown to calculate the kinematics and dynamics of obese participants. [22, 23] Research has also been carried out to determine the effects of removing thigh and shin markers on calculated kinematics [24, 25] as thigh and shin markers are generally more prone to soft tissue artefacts. [4, 7, 26]

This paper aims to test two hypotheses: a) The performance of MKO methods is adversely affected by body fat and b) time-based weighting [27] could be an effective way of reducing residual errors during the inverse kinematic step. The first hypothesis is tested by comparing residual error in the inverse kinematic step using three MKO methods on gait data collected using skin-based markers from people with varying body fat percentages as shown in Fig 1. Additionally, by altering the weight of the thigh marker at certain time instances through correlation of thigh-marker errors and total errors, we aim to test whether time-based weighting can lead to reduction in total residual error as shown in Fig 2. Thigh markers were chosen as they are significantly affected by STA. Improving the accuracy of MKO methods when applied to patients with varying levels of body fat, would aid the development of novel protocols that can more accurately analyse movement in a wider array of patients leading to more accurate diagnosis.

**Fig 1.**
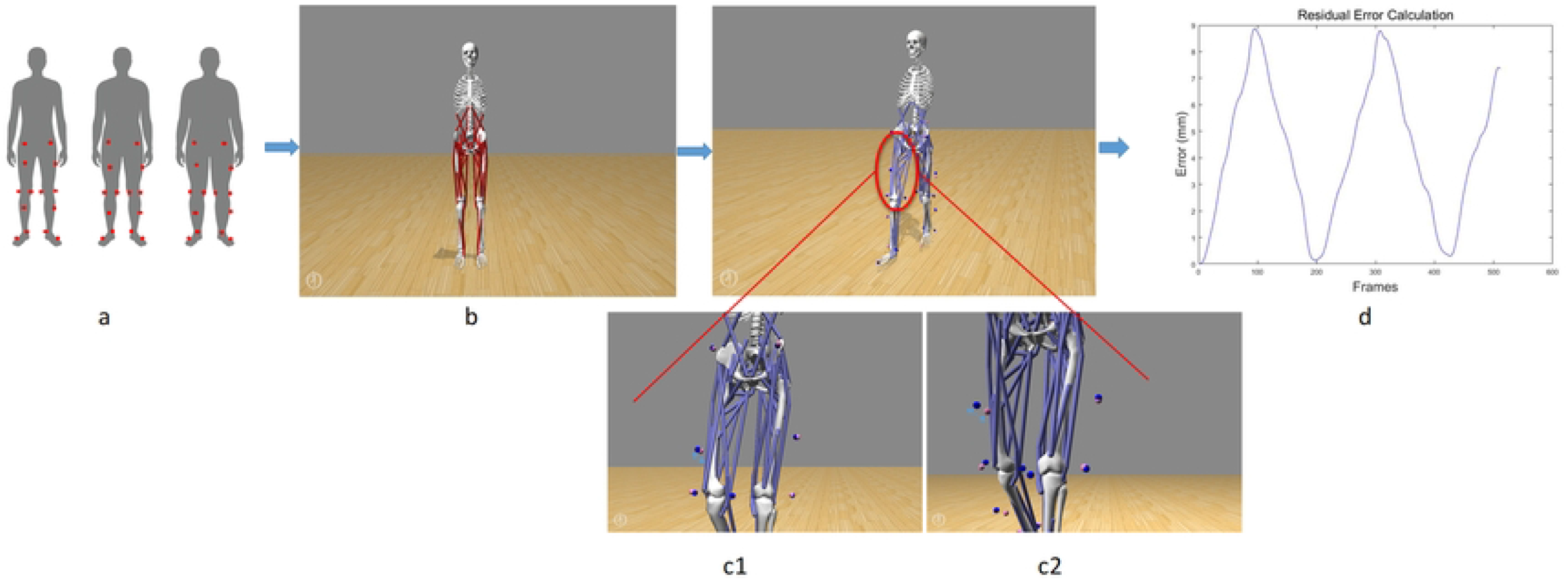
Workflow for analysing performance of MKO methods on varying body fat percentages. a) Collection of treadmill gait data using skin-mounted markers placed on participants of varying body fat. b) Generic OpenSim musculoskeletal model is scaled to subject specific dimensions using the gait data. c) Inverse kinematic analysis is performed using three different MKO methods. c1) Shows the difference in position of virtual marker (pink) and experimental marker (blue) for subjects of low body fat (blue arrow) and c2) shows the difference in virtual marker (pink marker) and experimental marker (blue marker) for subjects of high body fat (light blue arrow). d) Residual error obtained using three different MKO methods for subjects of low and high body fat is compared

**Fig 2.**
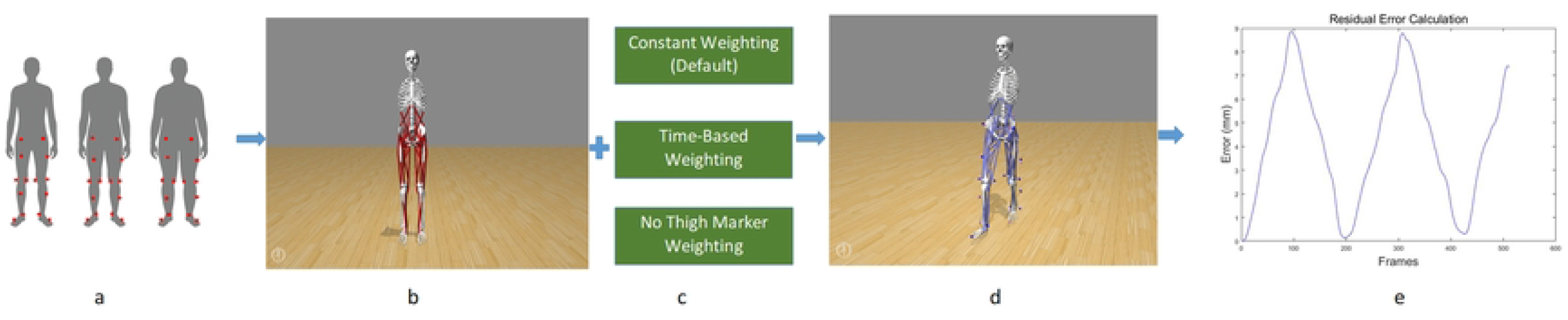
Workflow for analysing performance of time-based weighting for reducing residual error. a) Collection of treadmill gait data using skin-mounted markers placed on participants of varying body fat. b) Generic OpenSim musculoskeletal model is scaled to subject specific dimensions using the gait data. c) and d) Inverse kinematic analysis is performed using three different weighting schemes.e)Residual error obtained from the three different weighting scheme is compared

## Materials and methods

### Data collection

Skin-mounted marker based motion capture gait data was collected from 50 healthy participants of varying body fat percentage (*BMI*: 24.4 *±* 4). All the participants have provided written consent based on their understanding of the study, with ethics for the study provided by the Department of Electronics Engineering Ethics committee at the University of York, UK. Height and weight for each participant was recorded. Additionally, body fat at the thigh and hip were measured using skin calipers. Data was captured using Motive 2.4 and 10 Flex 3 cameras from OptiTrack.

Reflective markers were attached to each participant at specific anatomical landmarks on the lower body as specified by lower body Helen Hayes biomechanical markerset as shown in Fig 3. As part of the data collection protocol, participants were asked to stand stationary in a ‘T’ pose followed by walking on the treadmill at speeds of 0.1m/s, 0.3m/s and 0.5m/s. These speed values were determined beforehand in order to reflect standard clinical protocols. The raw data was filtered with a cut-off frequency of 10Hz; missing data frames were filled using either cubic interpolation function or pattern fill function, as provided in the software.

**Fig 3.**
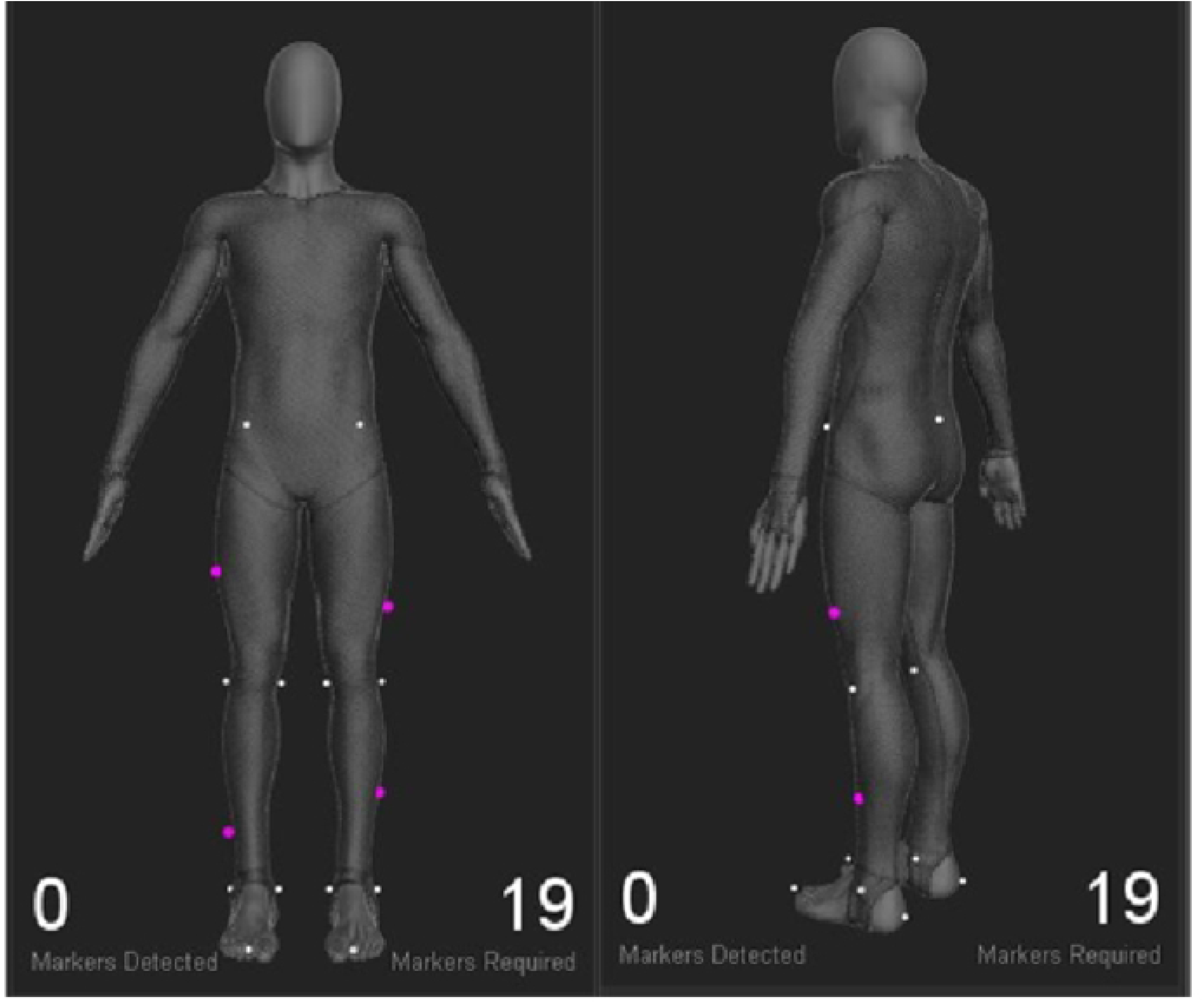
Helen hayes lower body biomechanics markerset.

### Residual error calculation

Residual errors from three MKO methods were calculated using musculoskeletal modelling software OpenSim. [28, 29] Residual error is calculated by taking the root mean square difference between the trajectory of the experimental marker and corresponding model marker in OpenSim. The three MKO methods analysed were the global optimisation method(GOM), [15] the Local marker estimation method [17, 30] and the Kalman smoothing method. [19] Prior to the calculation of residual errors using these three methods, OpenSim’s generic musculoskeletal model, gait2354, was scaled to subject-specific dimensions using either the static trial or the first frame of a dynamic trial. [31] Guidelines were followed to ensure that the error from the scaling procedure remained less than the value recommended by OpenSim. Based on the results of the inverse kinematic analysis, the total residual error for all the markers and marker-specific residual error was calculated for each participant at each walking speed.

To test the efficacy of time-based weighting scheme on reducing residual errors, a progressive weighting strategy was integrated into OpenSim; this weighting scheme has been previously applied to reduce errors resulting from occluded or missing markers. [27] The weight of a marker is progressively decreased to zero to coincide with the frame of interest, and progressively increased afterwards to its original value. The frame of interest and the weighting factor by which the weight is increased or decreased needs to be specified by the user. [27] The frame of interest was calculated by comparing the time-ranges of the peaks of total residual error and peaks of left and right thigh marker errors. Based on the number of gait cycles and walking speed, the number of frames of interest varies for each participant at each speed. Residual error using the time-based weighting scheme was compared to residual error calculated using the default weighting scheme, and to residual errors calculated by omitting the thigh markers. [24, 25] GOM was used as the MKO method for the inverse kinematic analysis in this case. Batch processing of OpenSim inverse kinematic analysis was performed in Python, whilst the residual error analysis was performed in MATLAB.

### Statistical analysis

Participant data was grouped based on both: thigh body fat measurements and body mass index (BMI) scores. As the majority of the participants were university students, the spread of body fat percentages and BMI scores was narrow. In order to perform satisfactory statistical analyses, participant data was grouped into six groups. Group 1 consisted of 15 participants with the highest body fat percentages. Group 2 consisted of 15 participants with the lowest body fat percentages. Group 3 and Group 4 consisted of 15 participants with the highest and lowest BMI scores respectively. Two additional groups, Groups 5 and 6, were created to account for variability in body fat measurements. Group 5 consisted of 10 participants with the highest body fat percentages which correlated with the highest BMI scores and Group 6 consisted of 10 participants with the lowest BMI scores which correlated with the lowest body fat percentages.

Statistical tests on total residual errors and right thigh residual error were performed for all the groups in pairs to test whether residual error calculated using the three MKO methods was affected by body fat. To determine the efficacy of time-based weighting, median values of total residual errors obtained using GOM and the three different weighting schemes were compared. Since not all Groups tested as normal using the Anderson-Darling test, Wilcoxon rank sum test was performed on all the unpaired data sets for testing whether MKO methods are adversely affected by higher body fat percentages. Wilcoxon signed rank test was performed on the paired data for testing the efficacy of time-based weighting on reducing residual errors. All statistical analyses were performed on MATLAB.

## Results

### MKO vs body fat percentages

Comparisons of total residual error for three different walking speeds obtained using GOM, LME and KS are shown in Fig 4, Fig 5, Fig 6 respectively. Box plots showing the difference in residual error at the thigh obtained using the three MKO methods for three different walking speeds is shown in Fig 7, Fig 8, Fig 9 respectively. As seen in the figures residual errors obtained using all three MKO methods was considerably larger for people with higher body fat percentages and BMI scores (Groups 1,3 and 5), compared to those with lower body fat percentages and BMI scores (Groups 2,4 and 6). This trend was observed for all walking speeds with residual error increasing at faster walking speeds for all participants. Total residual error was on average 30% higher in participants with higher body fat percentages and BMI scores, with residual error at the thigh on average higher by 25%.

**Fig 4.**
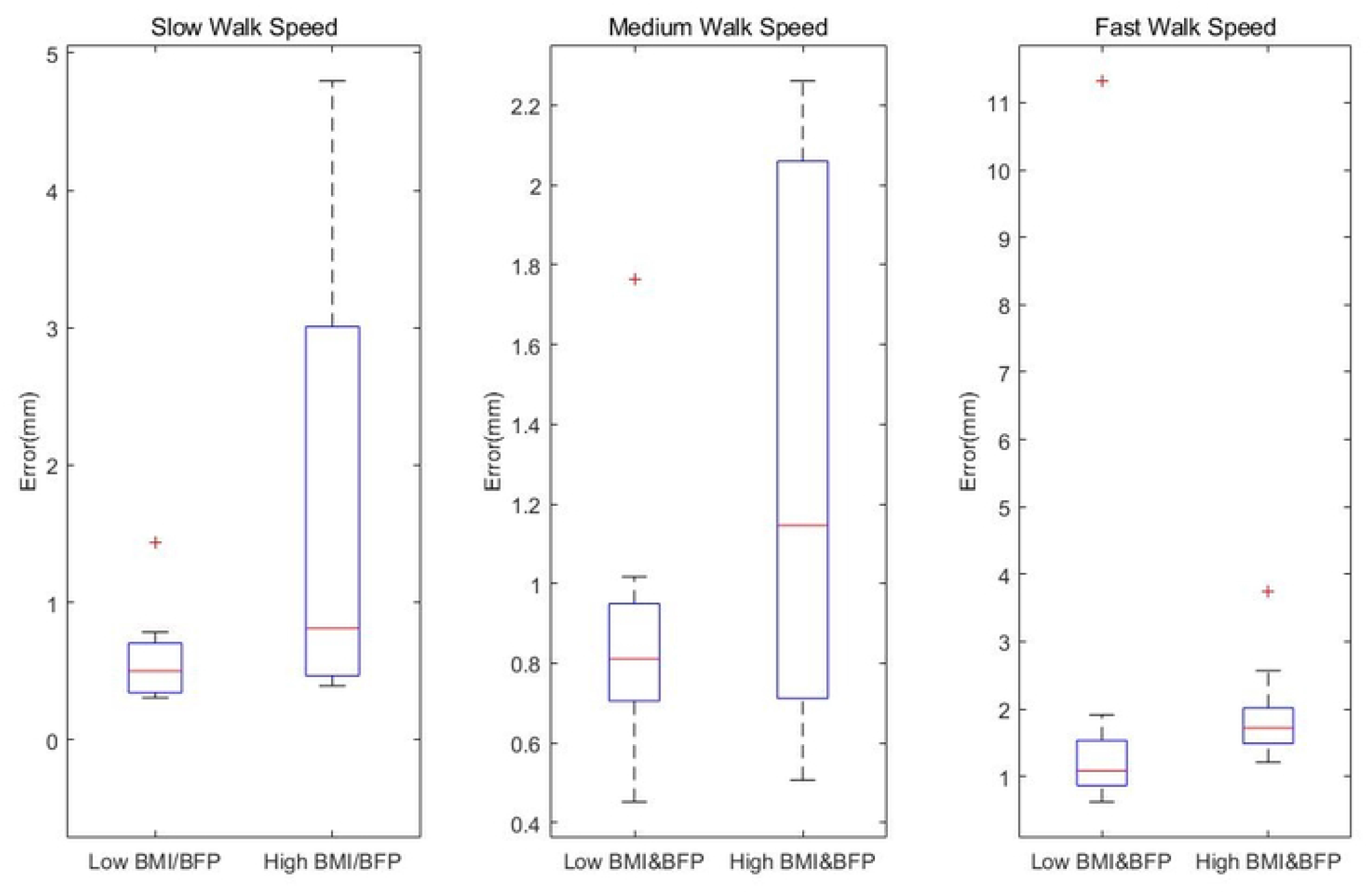
Total residual error comparison obtained using GOM. Comparison of total residual error between participants of low body fat percentages and high body fat percentages. a) Comparison at slow walking speed. b) Comparison at medium walking speed. c) Comparison at high walking speed.

**Fig 5.**
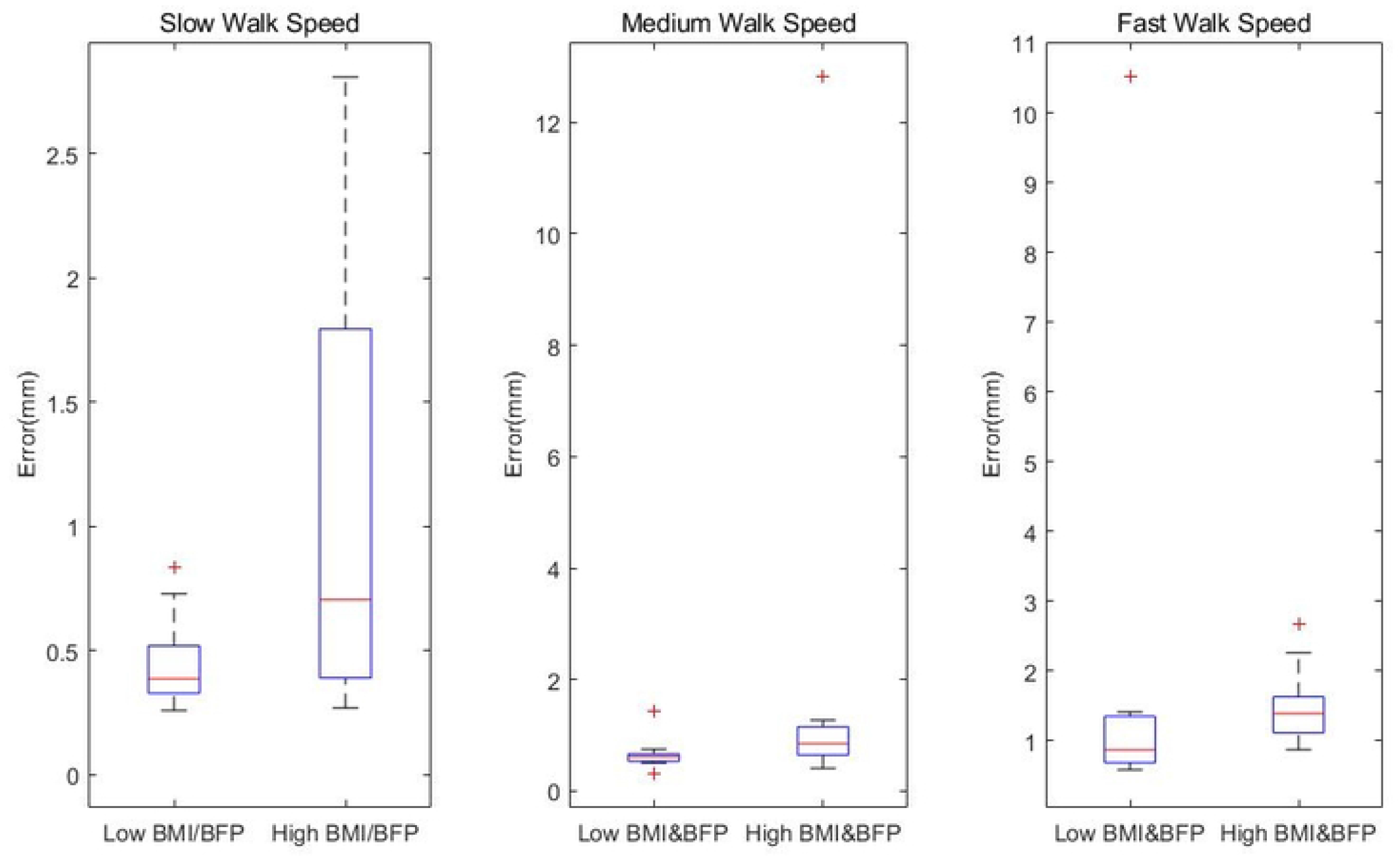
Total residual error comparison obtained using LME. Comparison of total residual error between participants of low body fat percentages and high body fat percentages. a) Comparison at slow walking speed. b) Comparison at medium walking speed. c) Comparison at high walking speed.

**Fig 6.**
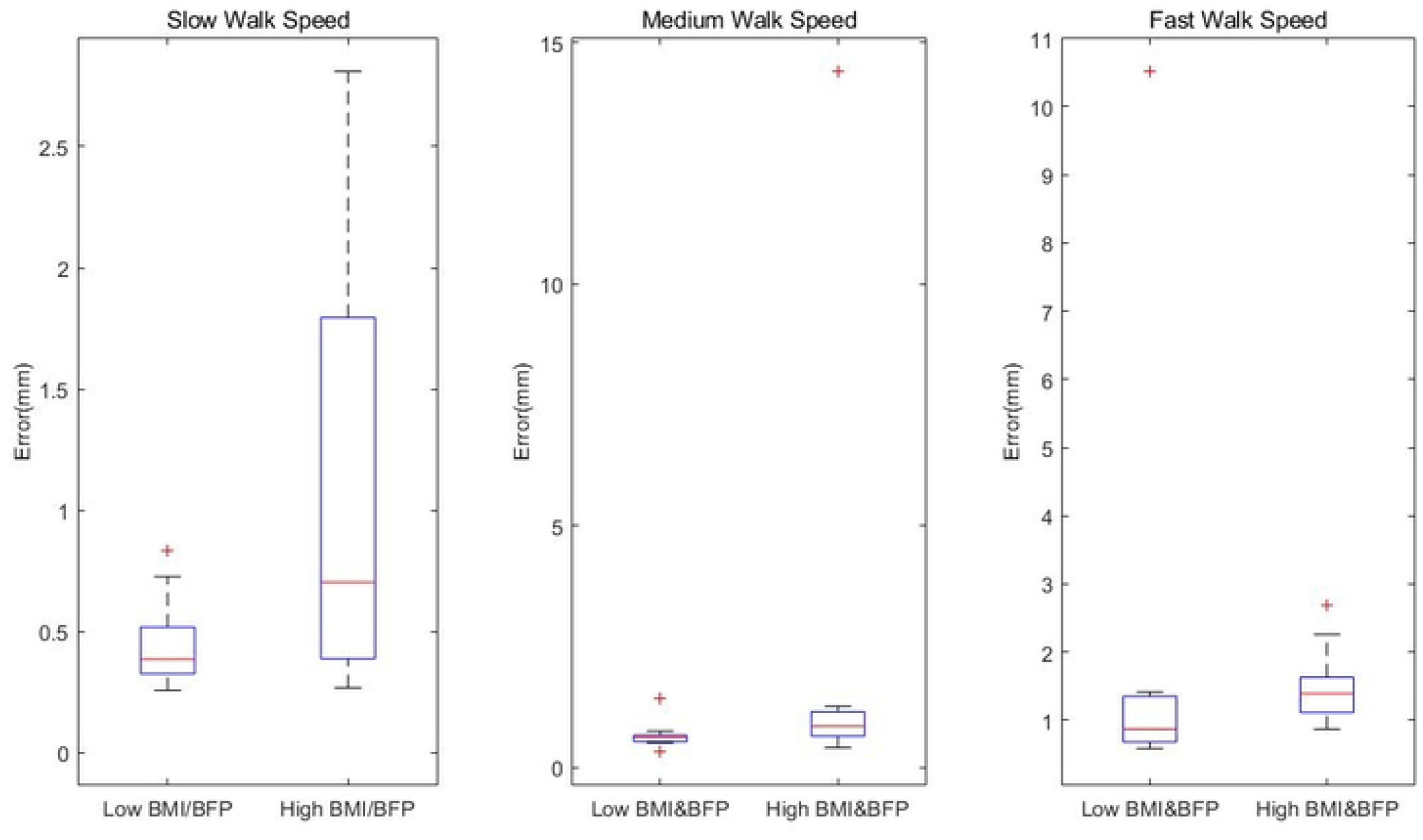
Total residual error comparison obtained using KS. Comparison of total residual error between participants of low body fat percentages and high body fat percentages. a) Comparison at slow walking speed. b) Comparison at medium walking speed. c) Comparison at high walking speed.

**Fig 7.**
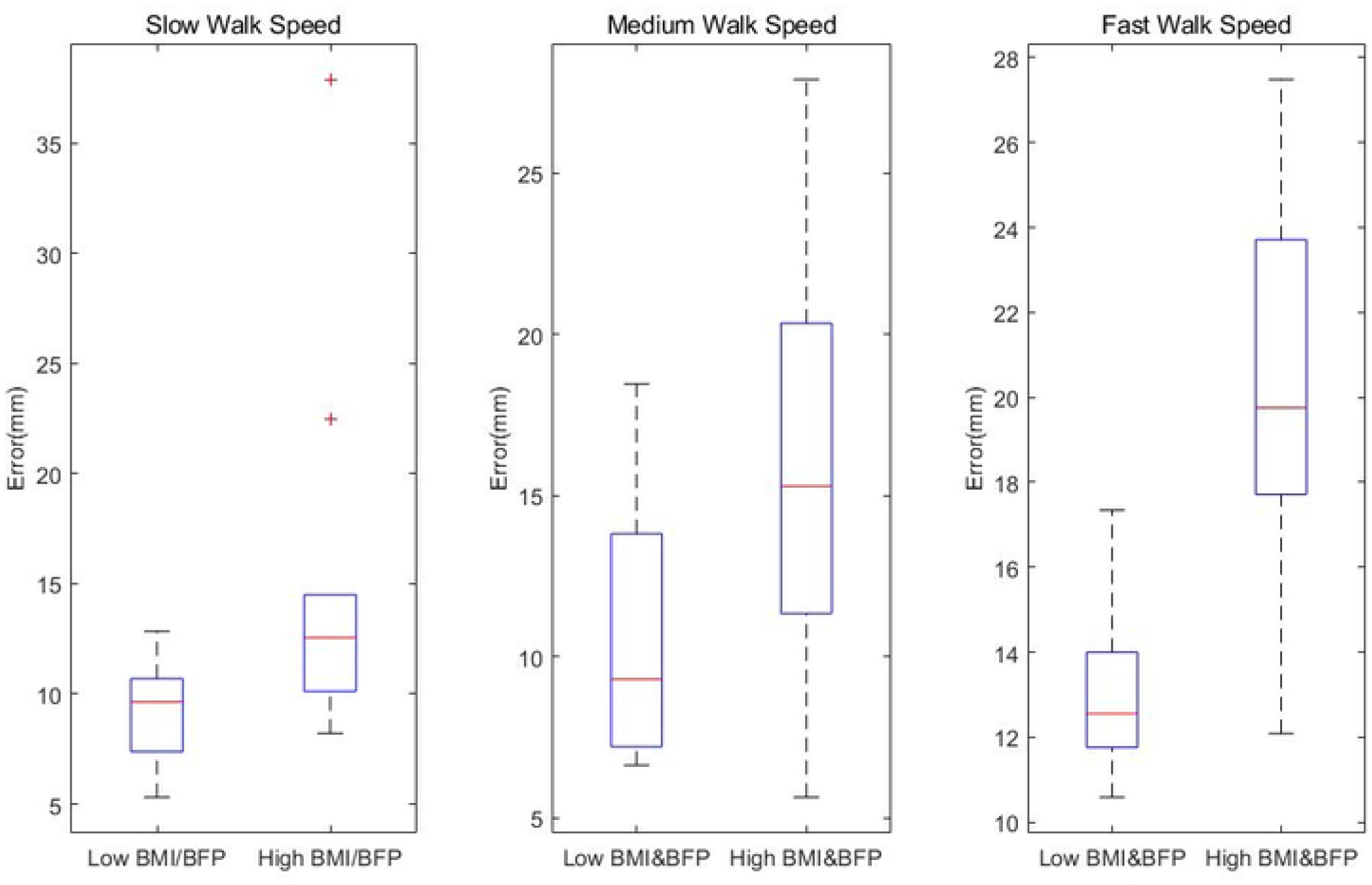
Thigh residual error comparison obtained using GOM. Comparison of residual error at thigh between participants of low body fat percentages and high body fat percentages. a) Comparison at slow walking speed. b) Comparison at medium walking speed. c) Comparison at high walking speed.

**Fig 8.**
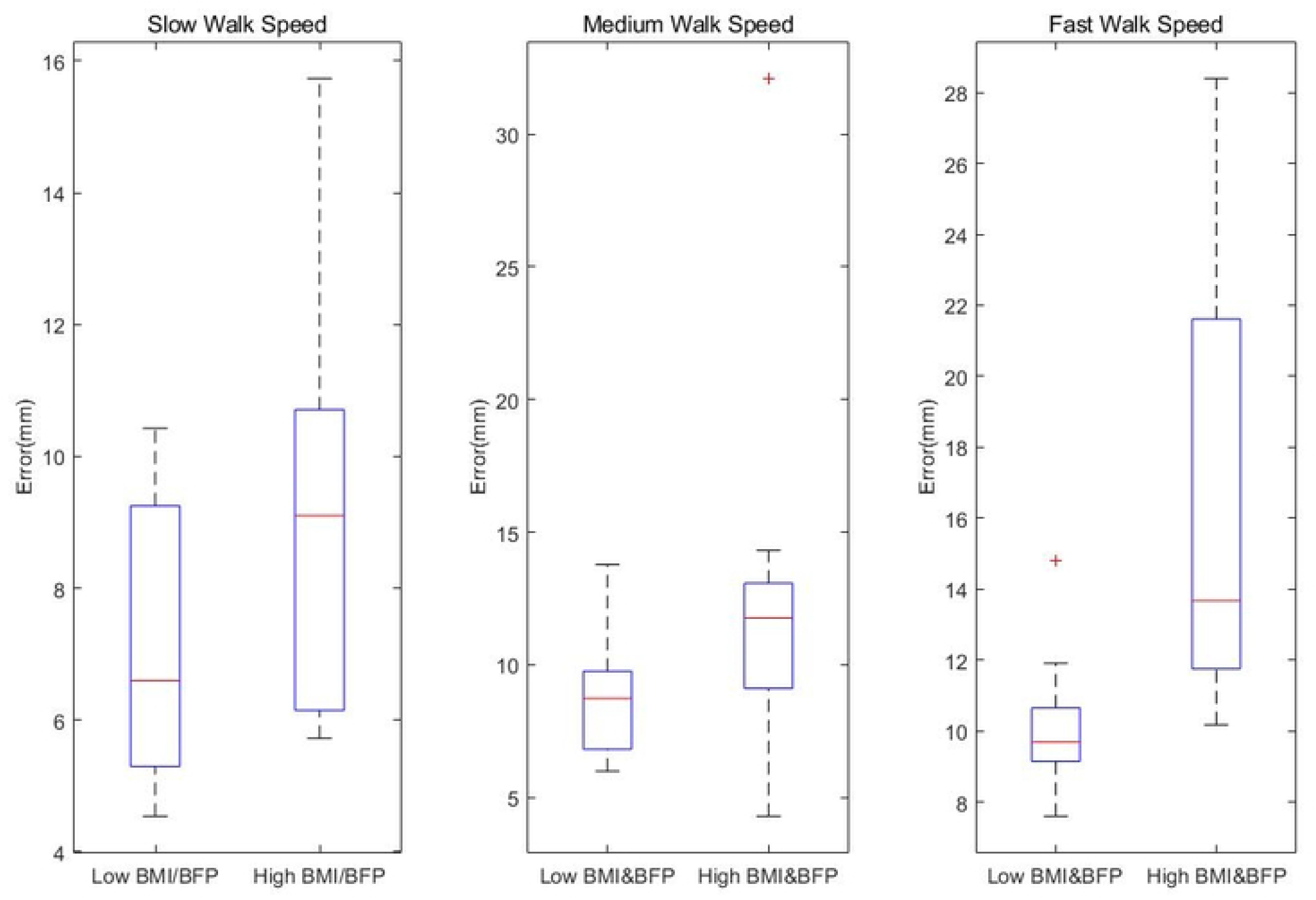
Thigh residual error comparison obtained using LME. Comparison of residual error at thigh between participants of low body fat percentages and high body fat percentages. a) Comparison at slow walking speed. b) Comparison at medium walking speed. c) Comparison at high walking speed.

**Fig 9.**
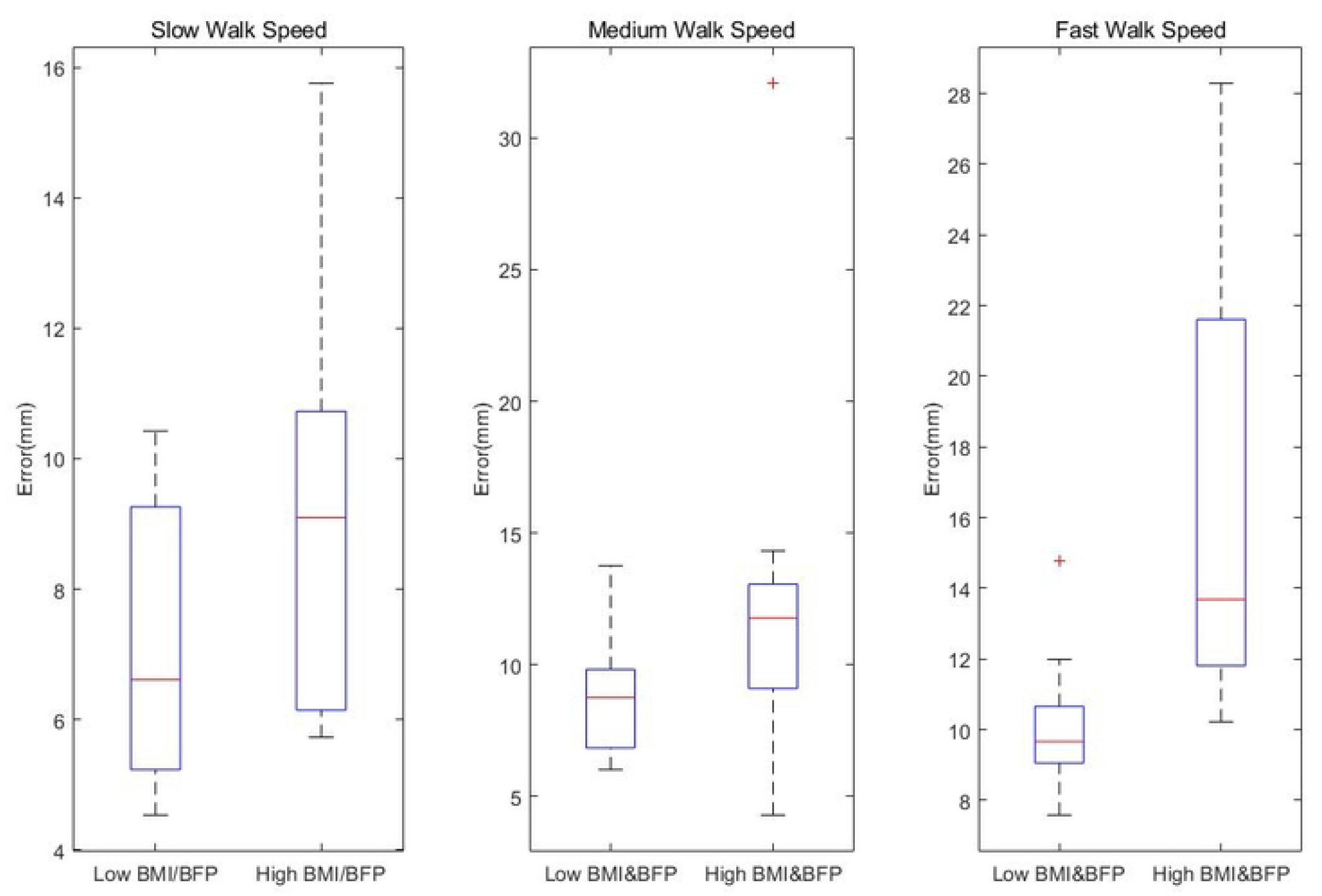
Thigh residual error comparison obtained using KS. Comparison of residual error at thigh between participants of low body fat percentages and high body fat percentages. a) Comparison at slow walking speed. b) Comparison at medium walking speed. c) Comparison at high walking speed.

The median values of total residual errors and median values of right thigh errors for Groups 1 and 2 are shown in Table1. As shown in Table X, median value of total residual error and right thigh error was greater for participants with high body fat percentages at all speeds and for all three MKO methods. At faster walking speeds, a Wilcoxon rank sum test indicated that the residual error at the thigh for Group 1 participants was significantly higher than residual error at the thigh for Group 2 participants when using local marker estimation (p=0.0421, Z=-2.03, N=15) and kalman smoothing (pp=0.0465, Z=-1.99, N=15). A Wilcoxon rank sum test on results obtained using GOM indicated that the difference in residual error at the thigh between Group 1 and Group 2 was statistically significant (high) at the fastest walking speed (p=0.0019, Z=-3.11, N=15).

**Table 1.** Median values of total residual error and residual error of thigh marker for Groups 1 and 2

| Method | Speed | Median marker errors |  |  |  |
| --- | --- | --- | --- | --- | --- |
|  |  | Body Fat Percentage (n=15) |  |  |  |
|  |  | Total Squared Error(m) |  | Right Thigh Error(m) |  |
|  |  | Low | High | Low | High |
| Global Optimisation Method | Slow | 0.00055 | 0.00082 | 0.01 | 0.014 |
|  | Intermediate | 0.00094 | 0.0012 | 0.0121 | 0.0147 |
|  | Fast | 0.0009 | 0.00139 | **0.0130\*\*** | **0.0179\*\*** |
|  | Overall | **0.0009\*\*** | **0.0013\*\*** | **0.0118\*\*** | **0.0155\*\*** |
| Local Marker Estimation | Slow | 0.00043 | 0.00068 | 0.0087 | 0.0095 |
|  | Intermediate | 0.00066 | 0.00097 | 0.00873 | 0.0121 |
|  | Fast | 0.001 | 0.0014 | **0.01\*** | **0.0134\*** |
|  | Overall | **0.00067\*** | **0.00098\*** | **0.0092\*** | **0.0117\*** |
| Kalman Smoothing | Slow | 0.00043 | 0.00068 | 0.0087 | 0.0095 |
|  | Intermediate | 0.00066 | 0.00097 | 0.00876 | 0.0122 |
|  | Fast | 0.001 | 0.0014 | **0.01\*** | **0.0135\*** |
|  | Overall | **0.00067\*** | **0.00098\*** | **0.0092\*** | **0.0119\*** |
| * : $p<0.05$ ** : $p<0.01$ *** : $p<0.001$ | | | | | |

Similar trends were observed in Groups 3 and 4 as shown in Table2. Wilcoxon rank sum test indicated that thigh residual errors for participants with higher BMI were significantly higher than thigh residual errors for participants with lower BMI obtained using local marker estimation (p=0.0279, Z=-2.53, N=15), kalman smoothing (p=0.0279, Z=-2.49, N=15) and GOM (p=0.0048, Z=-3.02, N=15).

**Table 2.**
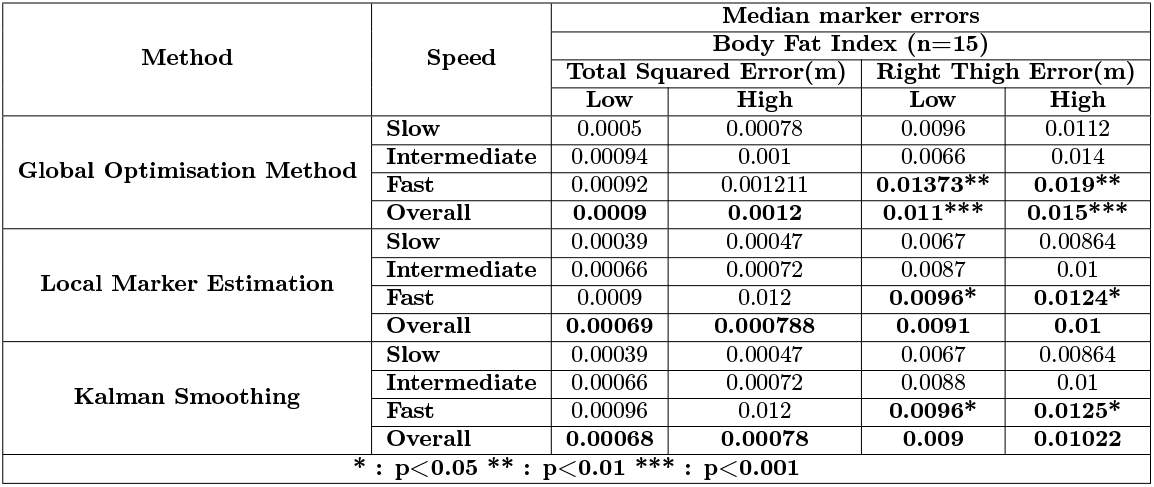
Median values of total residual error and residual error of thigh marker for Groups 3 and 4

| Method | Speed | Median marker errors |  |  |  |
| --- | --- | --- | --- | --- | --- |
|  |  | Body Fat Index (n=15) |  |  |  |
|  |  | Total Squared Error(m) |  | Right Thigh Error(m) |  |
|  |  | Low | High | Low | High |
| Global Optimisation Method | Slow | 0.0005 | 0.00078 | 0.0096 | 0.0112 |
|  | Intermediate | 0.00094 | 0.001 | 0.0066 | 0.014 |
|  | Fast | 0.00092 | 0.001211 | **0.01373\*\*** | **0.019\*\*** |
|  | Overall | **0.0009** | **0.0012** | **0.011\*\*\*** | **0.015\*\*\*** |
| Local Marker Estimation | Slow | 0.00039 | 0.00047 | 0.0067 | 0.00864 |
|  | Intermediate | 0.00066 | 0.00072 | 0.0087 | 0.01 |
|  | Fast | 0.0009 | 0.012 | **0.0096\*** | **0.0124\*** |
|  | Overall | **0.00069** | **0.000788** | **0.0091** | **0.01** |
| Kalman Smoothing | Slow | 0.00039 | 0.00047 | 0.0067 | 0.00864 |
|  | Intermediate | 0.00066 | 0.00072 | 0.0088 | 0.01 |
|  | Fast | 0.00096 | 0.012 | **0.0096\*** | **0.0125\*** |
|  | Overall | **0.00068** | **0.00078** | **0.009** | **0.01022** |
| * : p<0.05 ** : p<0.01 *** : p<0.001 |  |  |  |  |  |

The median values of total residual errors and median values of thigh residual error for Groups 5 and 6 are shown in Table3. Significant higher errors were observed in thigh residual errors of Group 5 when compared to Group 6 while using LME (p=0.0036,Z=-2.91,N=10), KS (p=0.0046,Z=-2.83,N=10) and GOM (p=0.001,Z=-3.28,N=10) at the fastest walking speed. Additionally, Wilcoxon rank sum test indicated an increase in total residual error that was statistically significant at the slowest walking speed (p=0.0452,Z=-2.00,N=10) and fastest walking speed (p=0.0036,Z=-2.15,N=10) when GOM was applied.

**Table 3.** Median values of total residual error and residual error of thigh marker for Groups 5 and 6

| Method | Speed | Median marker errors |  |  |  |
| --- | --- | --- | --- | --- | --- |
|  |  | Body Fat + Body Mass Index (n=10) |  |  |  |
|  |  | Total Squared Error(m) |  | Right Thigh Error(m) |  |
|  |  | Low | High | Low | High |
| Global Optimisation Method | Slow | **0.0005\*** | **0.0008\*** | **0.0096\*** | **0.01255\*** |
|  | Intermediate | 0.000811 | 0.0011 | 0.0092 | 0.0152 |
|  | Fast | **0.001\*** | **0.0017\*** | **0.01255\*\*\*** | **0.0197\*\*\*** |
|  | Overall | **0.00078\*\*** | **0.00144\*\*** | **0.0107\*\*\*** | **0.0164\*\*\*** |
| Local Marker Estimation | Slow | 0.00038 | 0.0007 | 0.0065 | 0.0091 |
|  | Intermediate | 0.0006 | 0.00084 | 0.0087 | 0.0117 |
|  | Fast | 0.00087 | 0.0013 | **0.0096\*\*** | **0.0136\*\*** |
|  | Overall | **0.0006\*\*** | **0.00097\*\*** | **0.0088\*\*** | **0.011194\*\*** |
| Kalman Smoothing | Slow | 0.00038 | 0.0007 | 0.00661 | 0.009 |
|  | Intermediate | 0.00062 | 0.000847 | 0.0087 | 0.01177 |
|  | Fast | 0.00087 | 0.00139 | **0.0096\*\*** | **0.0136\*\*** |
|  | Overall | **0.00063\*\*** | **0.00097\*\*** | **0.0088\*\*** | **0.01119\*\*** |
| * : p<0.05 ** : p<0.01 *** : p<0.001 |  |  |  |  |  |

Estimated marker trajectory for the thigh marker calculated using GOM, LME and KS for a participant with low body fat is shown in Fig 10,while the estimated trajectories calculated using the three methods for a participant for higher body fat percentage is shown in Fig 11. As seen in the figures, the estimated trajectories for the thigh marker obtained using the three MKO method matches the experimental marker trajectory for a participant with low body fat percentage but shows significant deviation for a participant with high body fat percentage.

**Fig 10.**
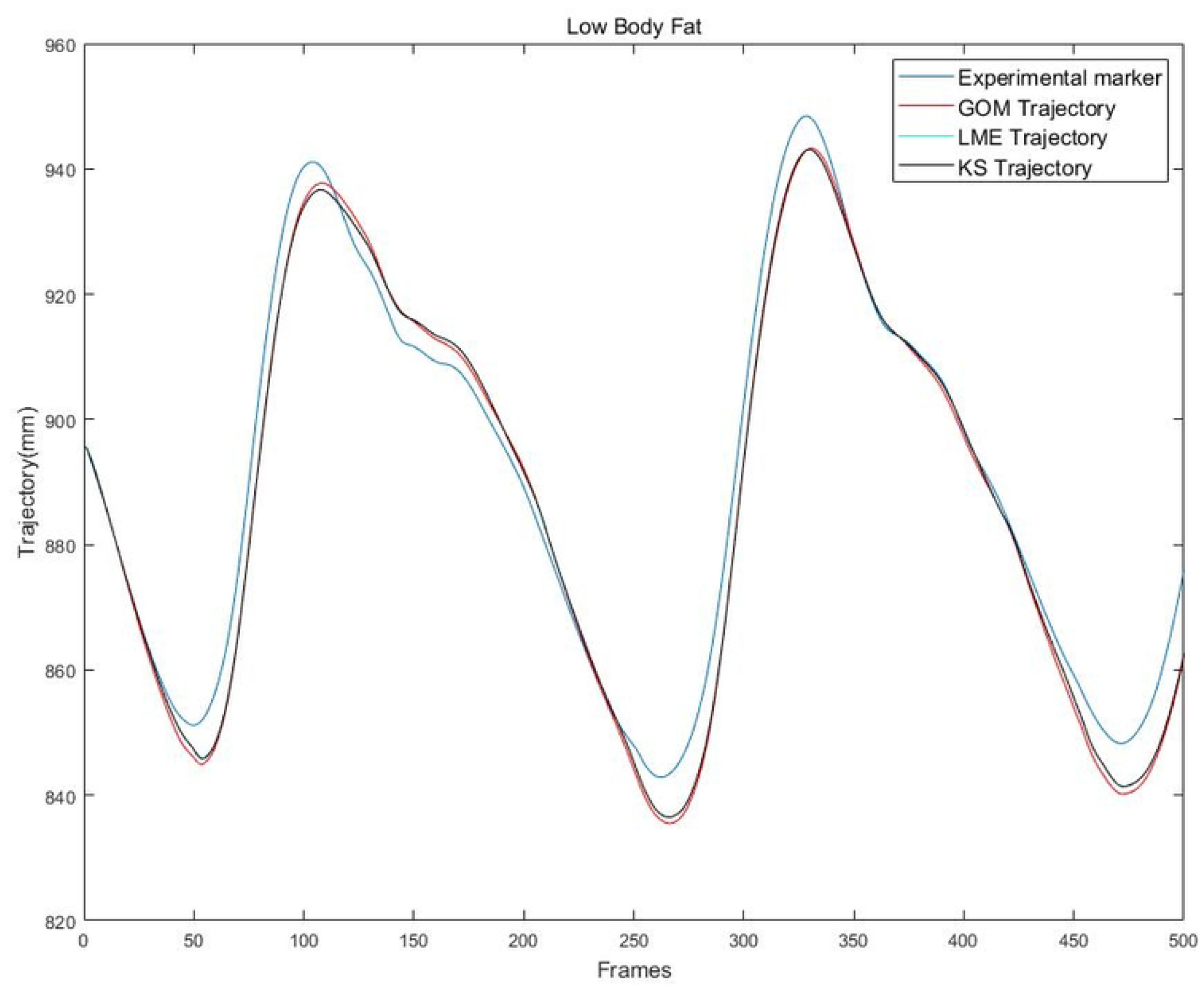
Comparison of estimated trajectories of thigh marker calculated using the three MKO methods and trajectory of experimental thigh marker for a participant of low body fat.

**Fig 11.**
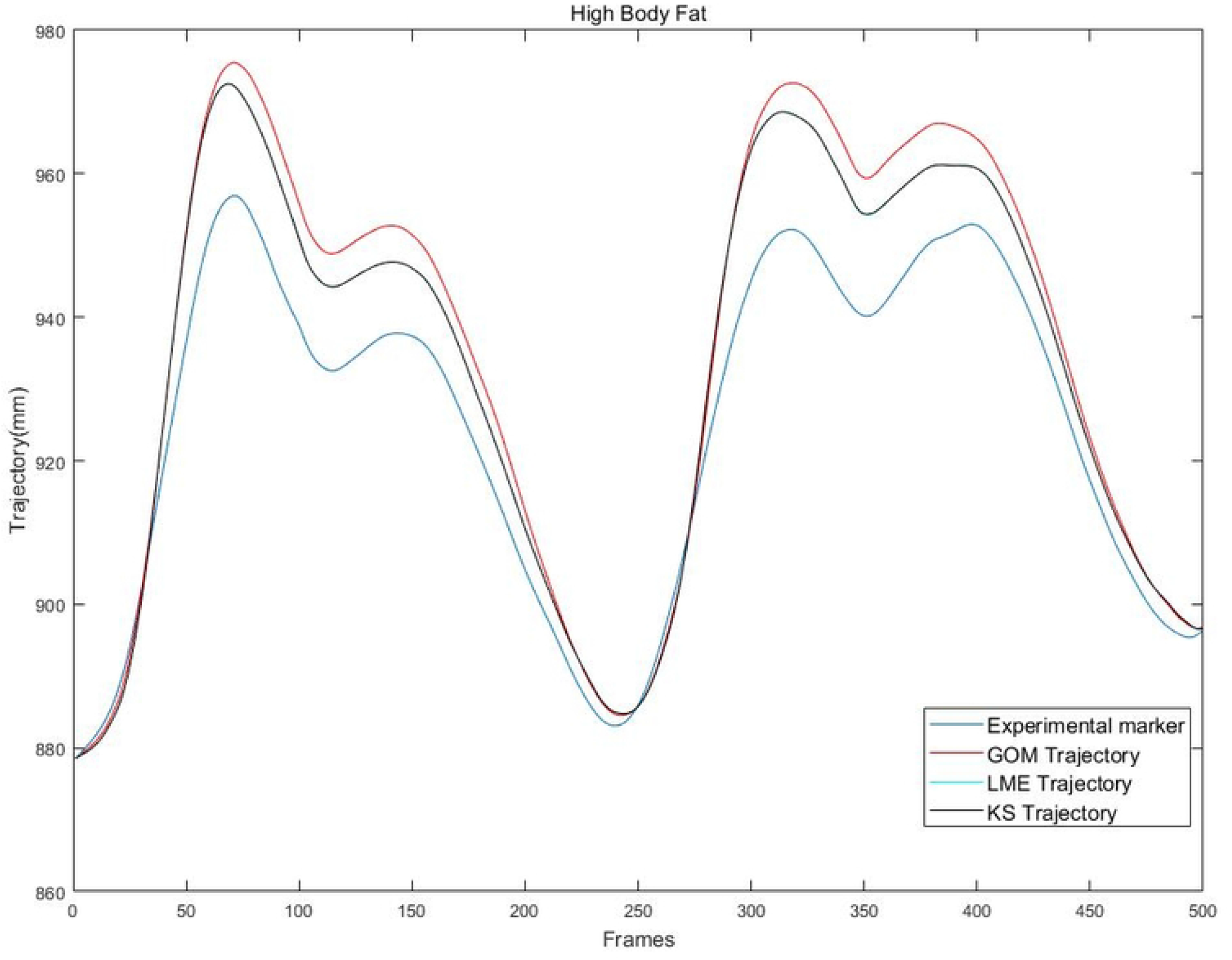
Comparison of estimated trajectories of thigh marker calculated using the three MKO methods and trajectory of experimental thigh marker for a participant of high body fat.

### Efficacy of time-based weighting

Application of time-based weighting showed a significant reduction in total residual errors compared to total residual errors obtained using the default weighting scheme and total residual errors obtained when thigh markers were omitted. Error reduction was maximum at the slowest walking speed and reduced at faster walking speeds.

Reduction in total residual error obtained using time-based weighting when compared to total residual error obtained using the default weighting scheme for the three walking speed was 58%, 54% and 40% for participants with higher body fat percentages and 55%, 51% and 29% for participants with lower body fat percentages. When compared to total residual error obtained when thigh markers were omitted, the reduction in error at the three walking speeds were 41%, 35% and 29% for participants with high body fat percentages and 29%, 29% and 14% for participants with low body fat percentages.A Wilcoxon signed rank test indicated that these results were statistically significant (p=0.0020,p=0.0020,p=0.0020,N=10) for high body fat percentages and low body fat percentages (p=0.0059,p=0.0020,p=0.0020,N=10).

Total residual error obtained using the three different weighting schemes for a participant with high body fat percentage are shown in Fig 12 while the total residual error obtained using the different weighting schemes for a participant with low body fat percentage is shown in Fig13. Variation of marker weight across frames calculated using time-based weighting is shown in Fig 14.

**Fig 12.**
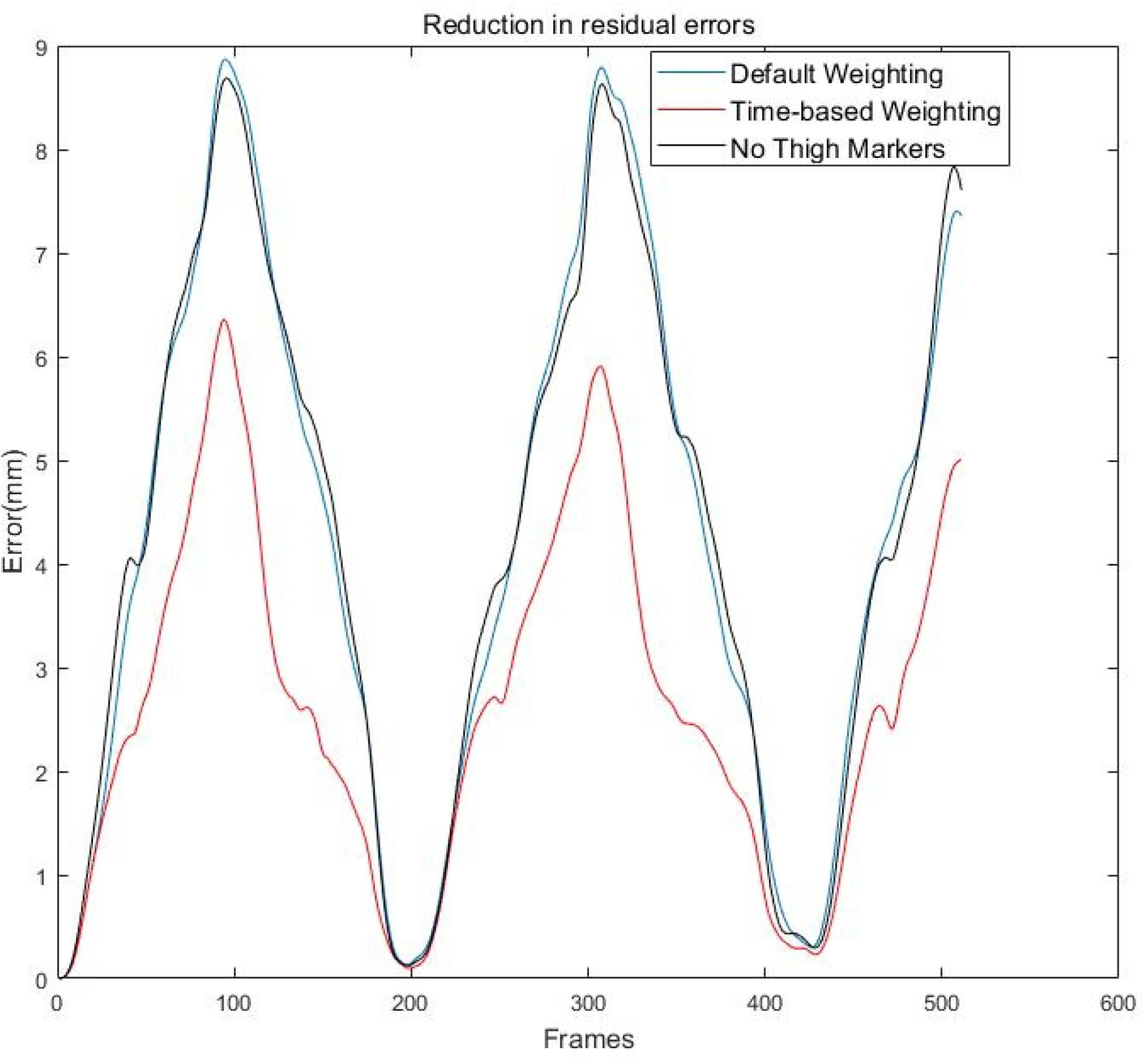
Comparison of total residual errors obtained using the three different weighting strategies for a participant of high body fat.

**Fig 13.**
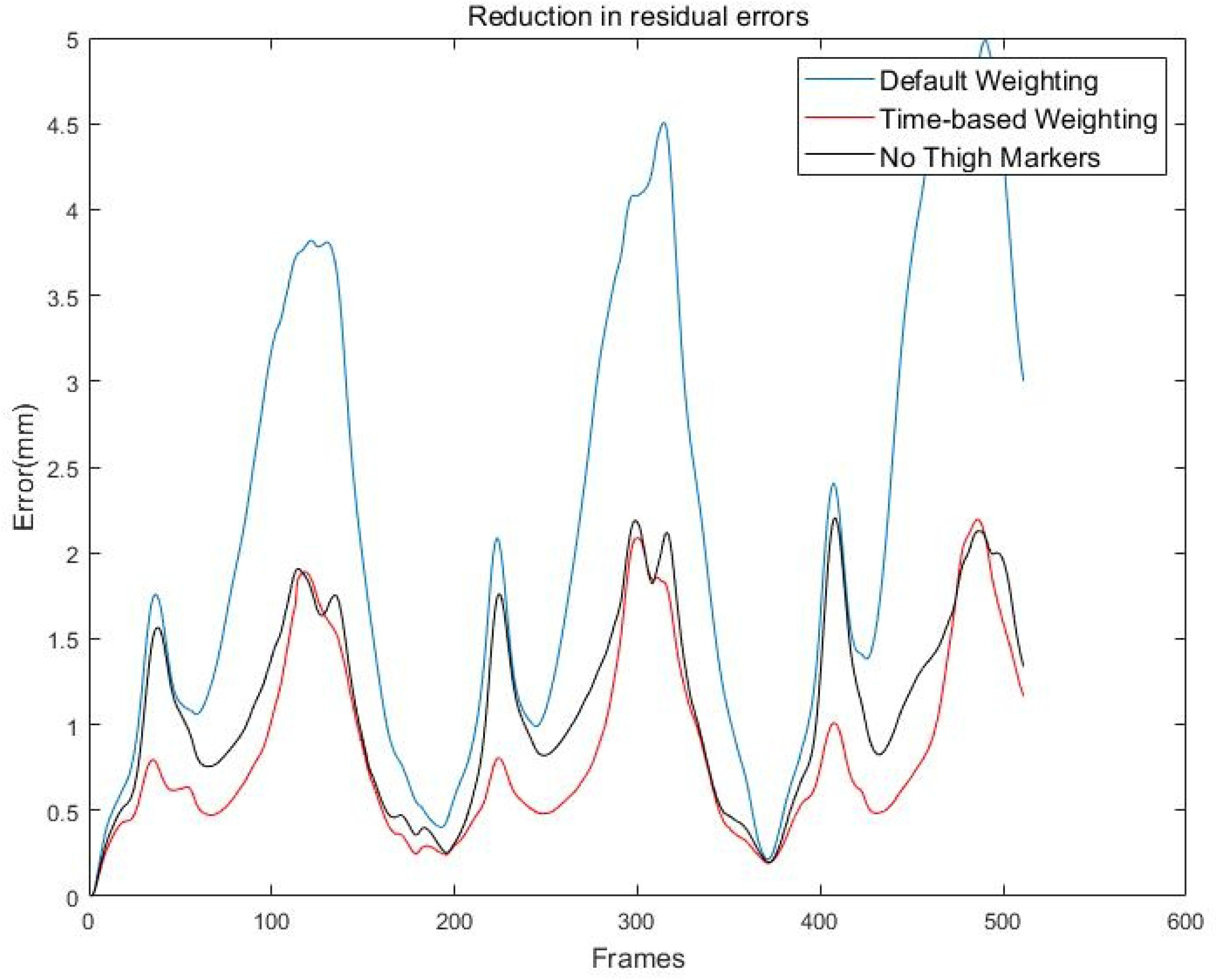
Comparison of total residual errors obtained using the three different weighting strategies for a participant of low body fat.

**Fig 14.**
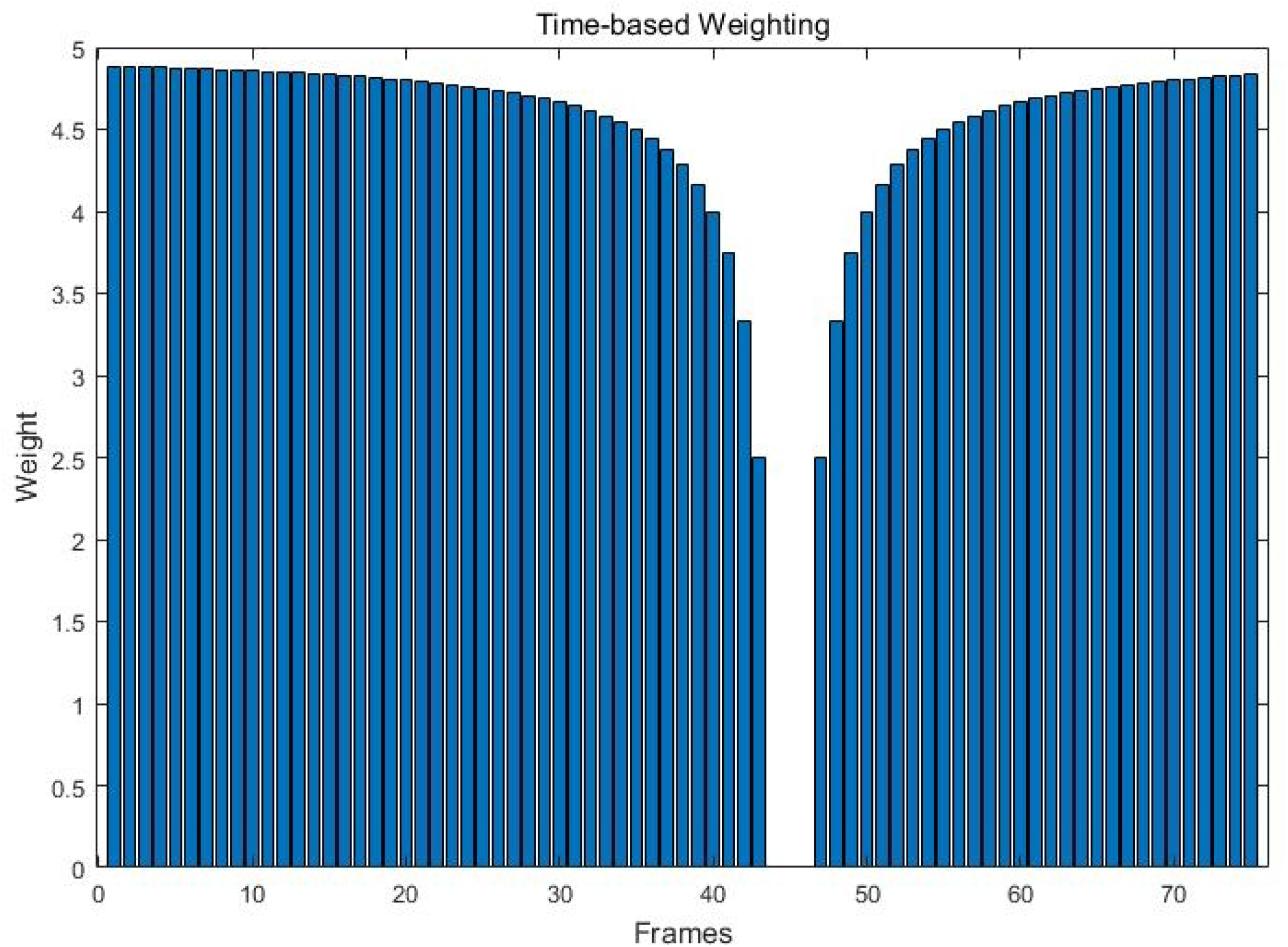
Time-based variation of marker weight. The marker weight is reduced as a combination of frames from reference frame and overall weight of the marker.

## Discussion

Soft tissue artefacts have been shown to adversely affect the clinical interpretation of marker based systems due to their influence on assessment of functional joint centres, and on the kinematic and dynamic analyses of various movement studies. [32, 33] Musculoskeletal modeling, through its incorporation of MKO methods for compensating soft tissue artefacts, has been increasingly used for movement analysis in both clinical and research settings. [10, 26] These optimisation methods [15, 19, 34, 35] have been predominantly validated on simulated movement data, with experimental data restricted to subjects with healthy or ideal body fat percentages and BMI scores. [7] Sensitivity analyses of musculoskeletal models to soft tissue artefacts have shown that soft tissue artefacts can cause variation of up to 30% in estimating joint forces. [8] These analyses, whilst indicating that soft tissue artefacts could adversely affect the results of kinematic analysis by magnitudes of five degree or greater, were performed on mathematical models of soft tissue artefacts [36] which do not account for body fat percentage or BMI scores in their formulation. In addition, to our knowledge, no analysis has been performed on the efficacy of MKO methods on movement data obtained from a large cohort of participants with varying body fat percentages.

Our primary hypothesis was that the efficacy of MKO methods is adversely affected by higher body fat percentages. Through the analyses and results, we have shown that residual errors obtained by three different MKO methods are significantly greater for the participants with higher body fat percentages and BMI scores compared to participants with lower body fat percentages and BMI scores. Inverse kinematics on data of the participants with higher body fat percentages had in general higher total residual errors by approximately 30% at every walking speed. Furthermore, residual errors at the right thigh were significantly higher at the fastest walking speed for all three methods. Whilst the residual errors were greater in higher body fat percentages compared to lower body fat percentages, the significant overlap between the ranges could be attributed to the exaggerated treadmill gait movement at very low speeds. GOM, which is the default MKO method applied in musculoskeletal modelling software OpenSim, showed the greatest differences in error when compared to kalman based methods such as KS and LME. This could be attributed to the fact that both the kalman based algorithms do not minimise errors based on a single frame, but work on temporal data, thereby making the trajectories smoother.

Our secondary aim was to verify whether time-based weighting could be used as an effective weighting scheme to reduce errors irrespective of body fat percentages when compared to OpenSim’s default weighting scheme. Additionally, we wanted to compare residual errors obtained using time-based weighting to residual errors obtained by excluding thigh markers. Research has shown that whilst inverse kinematics obtained while excluding thigh markers are highly correlated with results obtained when the all the markers were used, the greatest error reduction in joint angles was obtained when two markers were included in the thigh and shank. [24, 25, 37] Our results clearly demonstrate the efficacy of time-based weighting on reducing residual errors, compared to results obtained both by the default weighting scheme in OpenSim, and by excluding thigh markers. The reduction in error was highly significant (p<0.01) when comparing the results of time-based weighting to that of the default weighting scheme. The average reduction in error was 40% for participants with higher body fat percentages and 30% for participants of lower body fat percentages. Residual error was also reduced by 20% when thigh markers were not included for participants with higher body fat and by 10% for participants with lower body fat.

Through our analyses and results we have highlighted a major point of consideration while using musculoskeletal modelling for analysing biomechanics:that MKO methods, which are designed to reduce soft tissue artefacts, are adversely affected by higher body fat percentages and that additional measures are required when compensating for soft tissue artefacts resulting from varying body fat percentages. The amount and properties of soft tissues varies from individual to individual. This could result in varying magnitudes of soft tissue artefacts which are currently not compensated for in MKO methods. Analyses which have integrated wobbling masses into forward dynamic studies have shown that models integrating wobbling mass underestimate hip joint power by 50% and that as mass and inertia of the wobbling mass increases, the effects on joint kinematics becomes more extreme. [38] We have additionally shown that time-based weighting is an effective weighting strategy to reduce residual error compared to the default weighting schemes. We hypothesise that while precise methods to compensate for soft tissue artefacts which accounts for body fat percentages are being developed, awareness of the impact of body fat on MKO methods and integration of steps such as subject-specific scaling through MRI, incorporation of wobbling mass in inverse kinematics and weighting schemes such as time-based weighting can be an effective means of reducing soft tissue artefacts.

The main limitation of this study is that the analyses and results are based on comparisons of relative residual errors obtained from the inverse kinematic analysis in OpenSim and not from absolute residual errors obtained by comparing the results of inverse kinematic to that of ground truth data or true bone movement data. Thus the comparison of non-paired relative residual error or residual error of different individuals may affect the results underpinning the first hypothesis. Whilst all reasonable efforts have been made to ensure that each musculoskeletal model has been scaled to fit the individual of interest, the influence of different models on residual errors cannot be neglected. Future studies may look into incorporating the use of methods of ground truth data acquisition such as fluoroscopy or X-ray imaging to substantiate the above findings. While the above limitation has been alleviated in the analysis for the second hypothesis, time ranges for varying the weights were based on peaks of total residual errors and thigh errors, and not based on gait phases. An effective strategy for determining the time-ranges for reducing marker weights would be comparing how thigh residual errors impact the analysis during different gait phases and thereby reducing the weight based on the gait cycle. A limitation which could be alleviated by collecting data on a wider cohort of participants, is the lack of spread of body fat percentages and BMI scores resulting in reduced sample size for statistical analyses. Despite these limitations, this paper highlights the importance of accounting for variation in the performance of MKO methods based on body fat through different weighting schemes, and aims to lay the foundation upon which improvement in movement analysis using musculoskeletal modelling software for diagnosis and rehabilitation of musculoskeletal conditions can be achieved. We additionally believe our work can aid in the development of tools which can enhance our understanding and diagnosis of obesity related movement disorders. The prevalence of such disorders may vary based on ethnicity and socioeconomic conditions [39], therefore tools which address the effects of obesity on movement, may help to promote more equitable healthcare.

## Acknowledgments

We would like to thank the participants for their cooperation and involvement in the study. We would also like to thank OptiTrak for their technical support provided during the study.

## Author contributions statement

Conceptualisation of the idea, data collection and processing, analysis of results and writing of the original draft: Vignesh Radhakrishnan. Supervision, project administration, and editing of the paper: Samadhan B Patil. Supervision, and Review of the paper: Adar Pelah

